# Inhibition of *N*-myristoyltransferase Promotes Naive Pluripotency in Mouse and Human Pluripotent Stem Cells

**DOI:** 10.1101/2021.06.22.449326

**Authors:** Junko Yoshida, Hitomi Watanabe, Kaori Yamauchi, Takumi Nishikubo, Ayako Isotani, Satoshi Ohtsuka, Hitoshi Niwa, Hidenori Akutsu, Akihiro Umezawa, Hirofumi Suemori, Yasuhiro Takashima, Gen Kondoh, Junji Takeda, Kyoji Horie

## Abstract

Naive and primed states are distinct states of pluripotency during early embryonic development that can be captured and converted to each other *in vitro*. To elucidate the regulatory mechanism of pluripotency, we performed a recessive genetic screen of homozygous mutant mouse embryonic stem cells (mESCs) and found that suppression of *N*-myristoyltransferase (Nmt) promotes naive pluripotency. Disruption of *Nmt1* in mESCs conferred resistance to differentiation. Suppression of Nmt in mouse epiblast stem cells (mEpiSCs) promoted the conversion from the primed to the naive state. This effect was independent of Src, which is a major substrate of Nmt and is known to promote differentiation of mESCs. Suppression of Nmt in naive-state human induced pluripotent stem cells (hiPSCs) increased the expression of the naive-state marker. These results indicate that Nmt is a novel target for the regulation of naive pluripotency conserved between mice and humans.

## INTRODUCTION

Pluripotency is the potential to differentiate into three primary germ cell layers, and subsequently, adult tissues. Over recent decades, various approaches have been used to capture the pluripotent state in cell cultures^1^. Interestingly, these efforts have revealed that distinct pluripotent states can be established, both in mice and humans. The most widely studied pluripotent states are naive and primed states. The naive state represents a pluripotent state in pre-implantation-stage embryos from which mESCs are derived^2,3^, and the primed state corresponds to post-implantation-stage embryos, from which mEpiSCs are established^4,5^. In contrast, human embryonic stem cells (hESCs) were in the primed state under conventional culture conditions, despite being derived from pre-implantation embryos^6^. Human induced pluripotent stem cells (hiPSCs) were also in the primed state when established under the conventional hESC culture media^7,8^. Subsequently, naive-state hESCs/hiPSCs were established either by controlling signaling pathways with chemicals or through transient expression of transcription factors^9,10^. Elucidation of the mechanisms involved in regulating distinct states of pluripotency will provide clues for understanding the nature of pluripotency and its application in regenerative medicine.

We previously reported a method for the systematic generation of homozygous mutant mESCs, which entails the conversion of heterozygosity to homozygosity by transient inactivation of the Bloom’s syndrome gene (*Blm*)^11^. We extended this work and generated nearly 200 homozygous mutant mESC lines, especially for those genes with unknown functions. Through phenotypic screening of these homozygous mutant mESCs, we found that *Nmt1*-homozygous mutant mESCs are resistant to differentiation. Nmt is an enzyme that catalyzes the addition of a myristoyl group to the N-terminal region of proteins^12^. The inhibition of Nmt activity promoted the conversion of primed-state mEpiSCs into the naive state. Furthermore, the naive state of the hiPSCs was enhanced by an Nmt inhibitor, indicating that Nmt is an evolutionally conserved target for the regulation of naive pluripotency.

## RESULTS

### Disruption of Nmt1 Confers Differentiation Resistance to mESCs and Enhances the Properties of the Naive State

As one of the phenotypic screenings of our homozygous mutant mESC clones^11^, we sparsely plated each mESC clone on mouse embryonic fibroblasts (MEF) in serum-containing medium, obtained single cell-derived colonies, and assessed their morphology. Flat or small-sized colonies were occasionally observed in wild-type (*Wt*) mESCs (Fig. 1A). In contrast, *Nmt1*-homozygous mutant mESCs formed homogeneous colonies with round and dome shapes (Fig. 1A), suggesting that *Nmt1*-homozygous mutant mESCs are resistant to differentiation. To address this possibility, we maintained *Nmt1*-homozygous mutant mESCs in serum-containing medium without MEF feeder cells. From *Wt* mESCs, differentiated cells, such as enlarged cells with a decrease in the pluripotency marker Oct3/4, were prominent (Fig. 1B, left, arrowheads). In contrast, *Nmt1*-homozygous mutant mESCs formed tightly packed colonies that were Oct3/4-positive, even without MEFs (Fig. 1B, right), indicating a differentiation-resistant phenotype. To confirm that *Nmt1* mutation is responsible for this phenotype, we removed the gene trap vector sequence using Flp/*FRT* recombination to obtain revertant clones, using the protocol that we reported previously (Fig. 1C, top)^11^. The differentiation-resistant phenotype was abolished, as determined by the reduced number of alkaline phosphatase (ALP)-positive colonies (Fig. 1C, middle and bottom), demonstrating that *Nmt1* mutation is responsible for the differentiation-resistant phenotype. We also generated single cell-derived mESC colonies in serum-free N2B27 medium in the presence of 2i (inhibitors of MEK and GSK3) without LIF. The naive state of mESCs is stabilized under the serum-free 2i condition, which is called the ground state condition^13^. Both *Wt* mESCs and *Nmt1*-homozygous mutant mESCs formed tightly packed, undifferentiated colonies under the 2i condition (Fig. 1D). However, *Nmt1*-homozygous mutant mESCs were more dome-shaped than *Wt* mESCs (Fig. 1D). These results suggest that *Nmt1* deficiency not only confers differentiation resistance to mESCs but also promotes the naive state.

**Figure. 1.**
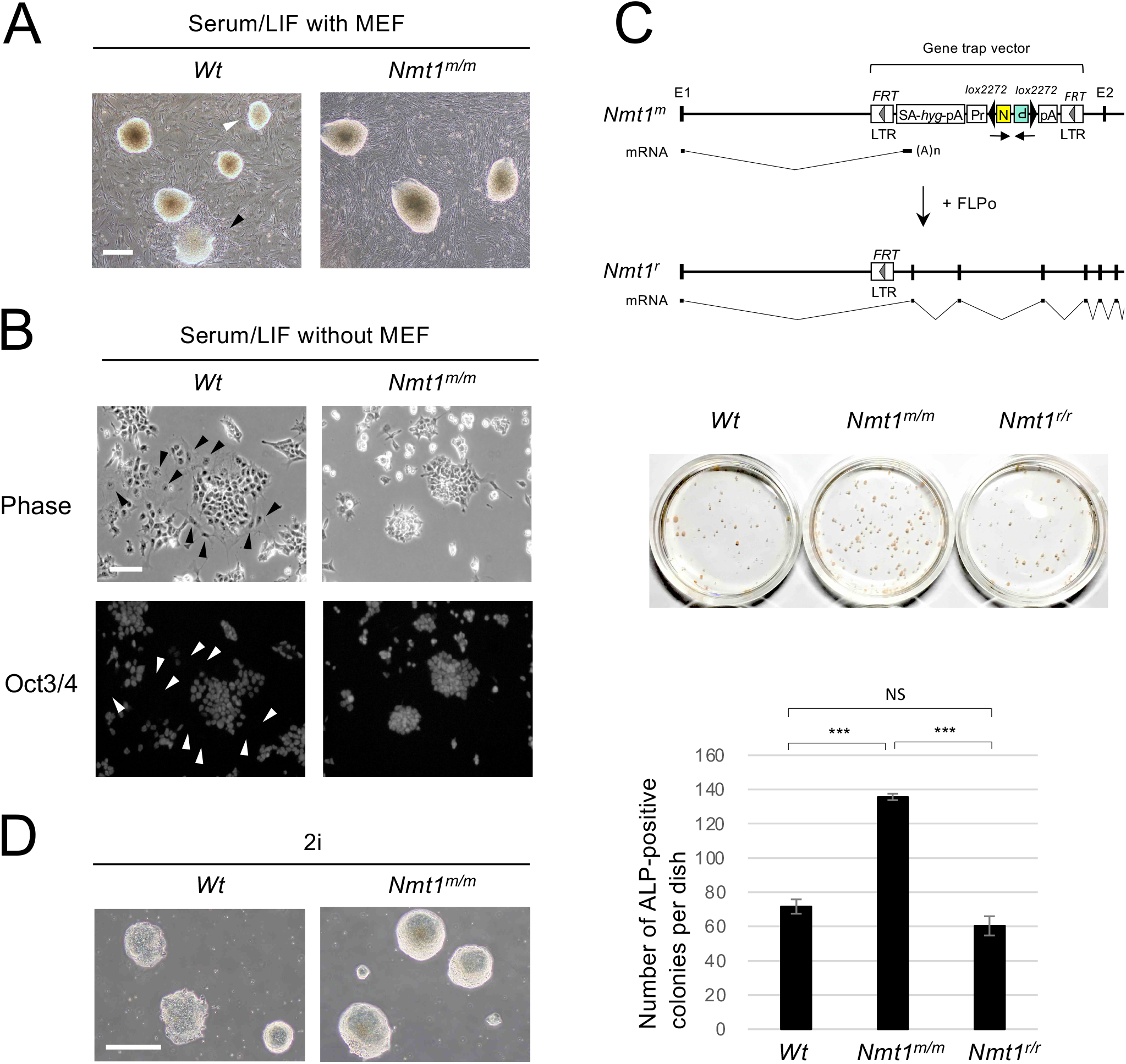
Disruption of Nmt1 confers differentiation resistance to mESCs and enhances properties of the naive state. (A) Morphological differences between wild-type (*Wt*) and *Nmt1*-homozygous mutant (*Nmt1*^*m/m*^) mESC colonies in serum/LIF medium. mESCs were sparsely plated on MEFs to obtain single cell-derived colonies. Flat (black arrowhead) or small-sized (white arrowhead) colonies were observed in *Wt* mESCs, whereas *Nmt1*^*m/m*^ mESCs were more homogeneous in shape and size and noticeably dome-shaped. Scale bar: 500 µm. (B) Oct3/4-staining of mESCs cultured for 12 days in serum/LIF medium without MEFs. *Nmt1*^*m/m*^ mESCs formed compact colonies with homogeneous Oct3/4-staining whereas *Wt* mESCs exhibited irregular-shaped colonies with scattered cells. Note that some *Wt* mESCs are enlarged and negative for Oct3/4 (arrowheads), indicating differentiation. Scale bar: 50 µm. (C) Reversion of the differentiation-resistant phenotype by deletion of the gene trap vector sequence. (Top) Generation of the revertant allele (*Nmt1*^*r*^) from the mutant allele (*Nmt1*^*m*^) by FLP/*FRT* recombination. (Middle, Bottom) Differentiation-resistant phenotype in each genotype. Six hundred mESCs were plated on MEFs in serum-containing medium without LIF, and the number of undifferentiated colonies was determined by ALP-staining. Data are shown as mean ± SEM (*n*=3, biological replicates). Tukey-Kramer test; ^***^ *p* < 0.001; NS > 0.05. E, exon; LTR, long terminal repeat; SA, splice acceptor; *hyg*, hygromycin-resistance gene; pA, polyadenylation signal; Pr, *Pgk1* promoter; N, neomycin-resistance gene; P, fusion gene of the puromycin-resistance gene and the herpes simplex virus thymidine kinase gene. Arrows below the gene trap vector indicates the orientation of each selection marker. (D) Morphological differences in single cell-derived colonies between *Wt* and *Nmt1*^*m/m*^ mESCs in serum-free 2i medium without LIF and MEFs. Note that *Nmt1*^*m/m*^ mESC colonies are noticeably dome-shaped compared to *Wt* mESC colonies. Scale bar: 500 µm.

### Conversion of Primed-state mEpiSCs into mESC-like Naive-state Cells by an Nmt Inhibitor

Differentiation resistance as well as the promotion of the naive state observed in *Nmt1*-homozygous mutant mESCs (Fig. 1) suggests that inhibition of Nmt1 activity may facilitate the conversion of primed-state mEpiSCs into mESC-like naive-state cells. To test this possibility, we used an Nmt inhibitor, DDD85646^14^. This inhibitor was originally reported as a lead compound against Nmt of *Trypanosoma brucei*, which causes African sleeping sickness. To test whether DDD85646 inhibits mammalian Nmt, we expressed an *N*-myristoylation signal-containing Venus (myrVenus) reporter^15^ in mESCs (Fig. 2A) and investigated the effect of DDD85646 on myrVenus localization (Fig. 2B). myrVenus was preferentially localized at cell membranes in the absence of DDD85646, and this preference was disrupted by DDD85646 (Fig. 2B), indicating that DDD85646 inhibits Nmt activity.

**Figure 2.**
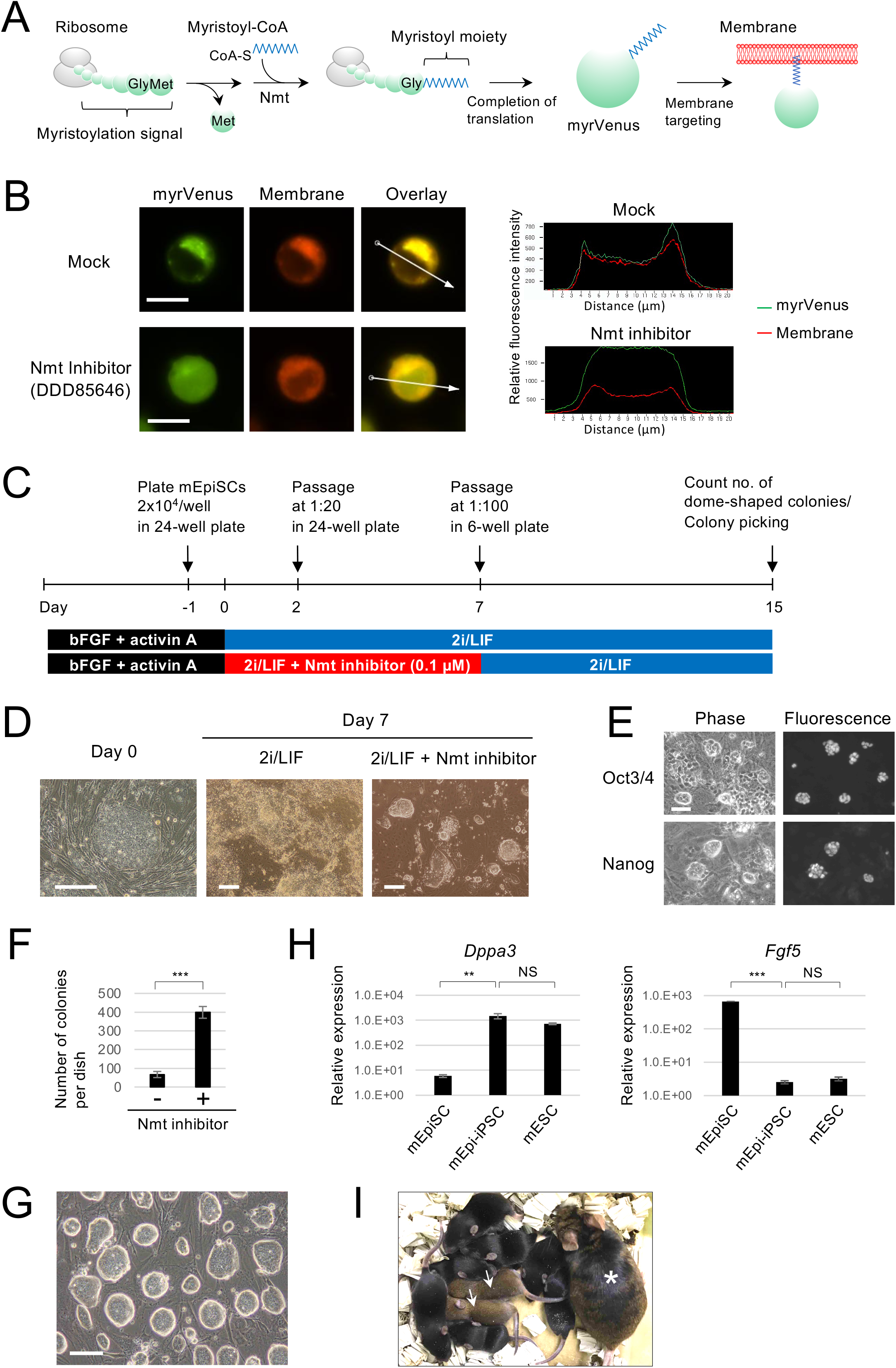
Conversion of primed-state mEpiSCs into mESC-like naive-state cells by the Nmt inhibitor. (A) Schematic of translation, myristoylation and membrane targeting of the *N*-myristoylation signal-containing Venus reporter (myrVenus). (B) The effect of the Nmt inhibitor DDD85646 on subcellular localization of myrVenus in mESCs. Line plots indicate the relative fluorescence intensity of myrVenus and membrane staining along the white arrow shown in the overlaid picture. The plasma membrane localization of myrVenus is decreased in the presence of the Nmt inhibitor. Scale bar: 10 µm. (C) Schematic of the protocol for the conversion of primed-state mEpiSCs into the mESC-like naive state. (D) Cells viewed at day 7 with or without the Nmt inhibitor. Dome-shaped mESC-like colonies were observed in the presence of the Nmt inhibitor. Scale bar: 200 µm. (E) Immunostaining of cells cultured under 2i/LIF + Nmt inhibitor. Dome-shaped colonies were positive for the pluripotency markers Oct3/4 and Nanog. Scale bar: 50 µm. (F) The number of dome-shaped colonies at day 15. Data are shown as mean ± SEM (*n*=3, biological replicates). Unpaired student’s *t*-test; ^***^ *p* < 0.001. (G) mESC-like cells stably maintained under 2i/LIF without the Nmt inhibitor. Scale bar: 500 µm. (H) mRNA expression of the naive-state marker *Dppa3* and the primed-state marker *Fgf5*. mEpi-iPSC indicates mESC-like cell induced from mEpiSC. Data are shown as mean ± SEM (*n*=3, biological replicates). Dunnett test; ^**^ *p* < 0.01, ^***^ *p* < 0.001, NS > 0.05. (I) Germline transmission of mEpi-iPSCs. An asterisk indicates a female parent chimera, and arrows denote agouti color-coated offspring.

Next, we tested whether the Nmt inhibitor facilitates the conversion of the primed-state mEpiSCs derived from post-implantation embryos into the mESC-like naive-state cells (Fig. 2C). There are distinct differences in the gene expression profile and growth conditions between mEpiSCs and mESCs. N2B27-based serum-free 2i medium supplemented with LIF (2i/LIF) is an optimal culture condition for naive-state mESCs, whereas this condition does not support primed-state mEpiSCs^4,5^. Consistent with this idea, high levels of cell death and differentiation were observed in mEpiSCs under 2i/LIF (Fig. 2D, middle). A combination of 2i/LIF and the Nmt inhibitor DDD85646 also induced cell death and differentiation; however, dome-shaped mESC-like colonies appeared after one week (Fig. 2D, right). These dome-shaped colonies were positive for the pluripotency markers Oct3/4 and Nanog (Fig. 2E), suggesting that mEpiSCs were successfully converted into the mESC-like naive state cells. After replating, the cells were cultured under 2i/LIF without the Nmt inhibitor. The number of dome-shaped colonies was substantially greater when the cells were pretreated with the Nmt inhibitor (Fig. 2F). After picking each dome-shaped colony, we could establish mESC-like clones under 2i/LIF without the Nmt inhibitor (Fig. 2G). The gene expression pattern of these clones, named mEpi-iPSC clones, was similar to that of mESCs, with high levels of expression of the naive-state marker *Dppa3* and low levels of expression of the primed-state marker *Fgf5* (Fig. 2H), strongly suggesting that mEpi-iPSC clones were in the naive state. To further demonstrate the conversion to the naive state, we injected each mEpi-iPSC clone into pre-implantation mouse embryos (8-cell stage embryos or blastocysts) and generated chimeric mice. The efficiency of generating chimeric mice is extremely low when using mEpiSCs^4,5^; however, we successfully generated chimeric mice from six out of seven mEpi-iPSC clones (Fig. 2I; Supplementary Fig. 1). Furthermore, we could achieve germline transmission in four clones (Fig. 2I; Supplementary Fig. 1), confirming that these clones were in the naive state. These results indicate that the suppression of Nmt promotes the conversion of the primed state into the naive state.

### Validation of the Effect of Nmt1 Deficiency on the Primed to Naive Conversion by Conditional *Nmt1* Knockout

To confirm that the effect of the Nmt inhibitor DDD85646 on the primed to naive conversion was not the off-target effect but the on-target effect, we genetically inactivated the *Nmt1* gene in the primed state and examined whether *Nmt1*-inactivation induces a naive state, as outlined in Fig. 3A. First, we manipulated the *Wt* allele of the *Nmt1*-mutant heterozygous mESC line and generated the floxed *Nmt1* allele (Fig. 3A; Supplementary Fig. 2). The parental mESC line of this mutant contains the *ERT2-iCre-ERT2* fusion recombinase gene at the *Rosa26* locus^11,16^; therefore, the floxed *Nmt1* allele can be conditionally inactivated by 4-hydroxytamoxifen (4HT). Next, we induced the *Nmt1*-floxed mESC line into the mEpiSC-like primed-state cell line using bFGF and activin A, according to the published protocol^17^ (Fig. 3A). Last, we inactivated Nmt1 in the primed state using 4HT and cultured under 2i/LIF to examine whether Nmt1-deficiency induces conversion of the primed state into the naive state. Deletion of *Nmt1* was confirmed by PCR analysis of the *Nmt1* locus (Fig. 3B) and reduced membrane localization of the myrVenus reporter (Fig. 3C). The elimination of membrane localization was not complete (Fig. 3C). There are two *Nmt* genes in mice, *Nmt1* and *Nmt2*, and Nmt2 is expressed in mouse blastocysts although the expression level is lower than Nmt1^18^. We speculate that the residual membrane localization of the myrVenus reporter (Fig. 3C) is due to Nmt2 activity.

**Figure 3.**
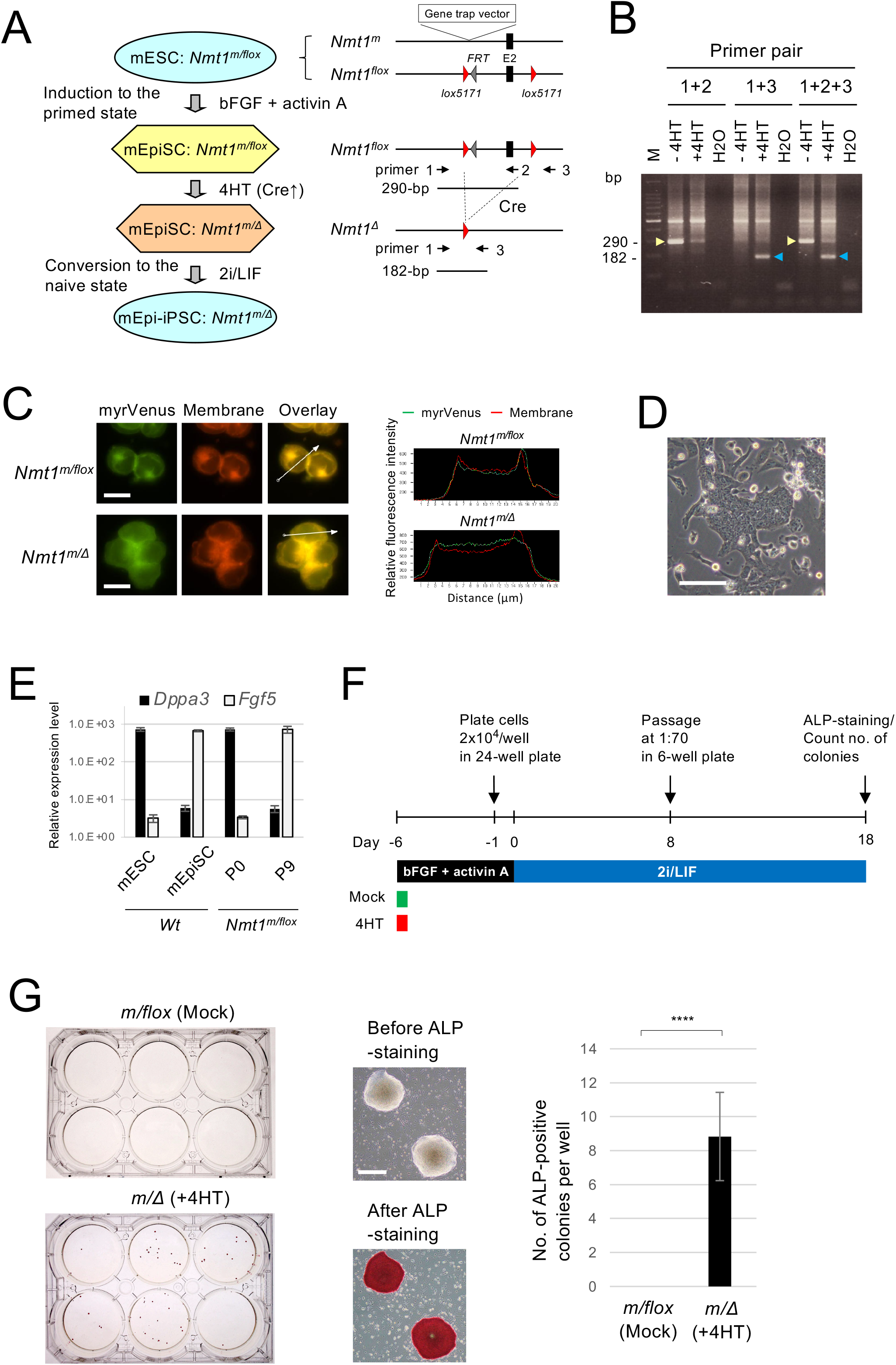
Verification of the effect of Nmt1 deficiency by conditional gene knockout. (A) Outline of the generation of genetically modified mEpiSC-like cells and conversion to naive-state mEpi-iPSCs by conditional knockout of Nmt1. 4HT, 4-hydroxytamoxifen. (B) PCR analysis of the 4HT-induced deletion of the *Nmt1* gene. Primer pairs are depicted in (A). PCR bands derived from the *Nmt1*-floxed allele and the 4HT-induced deleted allele are indicated by yellow and blue arrowheads, respectively. M, 100-bp size marker. (C) Effect of *Nmt1* knockout on subcellular localization of the myrVenus reporter. Line plots indicate the relative fluorescence intensity of myrVenus and membrane staining along the white arrows shown in the overlaid picture. Plasma membrane localization is decreased by 4HT-induced conditional knockout. Scale bar: 10 µm. (D) A morphological view of mEpiSC-like cells induced from mESCs by continuous culturing under bFGF and activin A at passage 9. Scale bar: 100 µm. (E) Confirmation of the acquisition of the primed-state features in mEpiSC-like cells shown in (D). P0, passage 0; P9, passage 9. (F) Schematic of the protocol for the conversion of primed-state mEpiSC-like cells into the naive-state mEpi-iPSCs. (G) (Left) ALP-staining of the colonies after the conversion process. (Middle) Examples of ALP-positive colonies before and after ALP-staining. Scale bar: 500 µm. (Right) Number of ALP-positive colonies are shown as mean ± SEM (*n*=3, biological replicates). Note that no ALP-positive colony was obtained in mock-treatment. Unpaired student’s *t*-test; ^****^ *p* < 0.001.

After continuous culture of mESCs under bFGF and activin A, we obtained mEpiSC-like flat colonies (Fig. 3D). mEpiSC-like features were confirmed by a decrease in the naive-state marker *Dppa3* and the induction of the primed-state marker *Fgf5* (Fig. 3E). We then inactivated the *Nmt1* gene with 4HT and induced conversion of the primed state to the naive state under 2i/LIF (Fig. 3F). ALP-positive dome-shaped colonies appeared by inactivating Nmt1, whereas no ALP-positive colonies were obtained in mock-treatment (Fig. 3G). The results were consistent with the observation in the Nmt inhibitor (Fig. 2F), demonstrating that Nmt1 deficiency promotes conversion of the primed state to the naive state.

### The Effect of the Nmt Inhibitor on the Primed to Naive Conversion Is Not Mediated by Src Signaling Pathways

Next, we searched for the Nmt substrate associated with the regulation of naive pluripotency. Many proteins have been reported as substrates of Nmt^12^. Among them, we focused on Src for the following reasons. First, a Src inhibitor supports the maintenance of the naive state in mESCs and can replace the MEK inhibitor in serum-free 2i/LIF culture^19^. Second, a Src inhibitor was included in a chemical cocktail for establishing naive hESCs/hiPSCs^10^. Therefore, we considered that the effect of the Nmt inhibitor observed in our study could be mediated through the inhibition of Src kinase activity.

To test this possibility, we treated mEpiSCs with the Src inhibitor CGP77675^19^ and compared the effect on primed to naive conversion with that of the Nmt inhibitor DDD85646 (Fig. 4A). According to the previous report^19^, the optimal concentration of CGP77675 for maintaining the naive state in mESCs is 1.5 µM. Therefore, we tested a wide range of concentrations, from 0.5 to 6 µM, including the optimal concentration for mESCs (Figs. 4A and 4B). However, we observed almost no increase in the efficiency of primed to naive conversion (Fig. 4C). At high-range concentrations (≥4 µM), we simply observed severe growth suppression (Fig. 4B). These results indicate that unknown factors other than Src kinase are responsible for the effect of the Nmt inhibitor on the primed to naive conversion.

**Figure 4.**
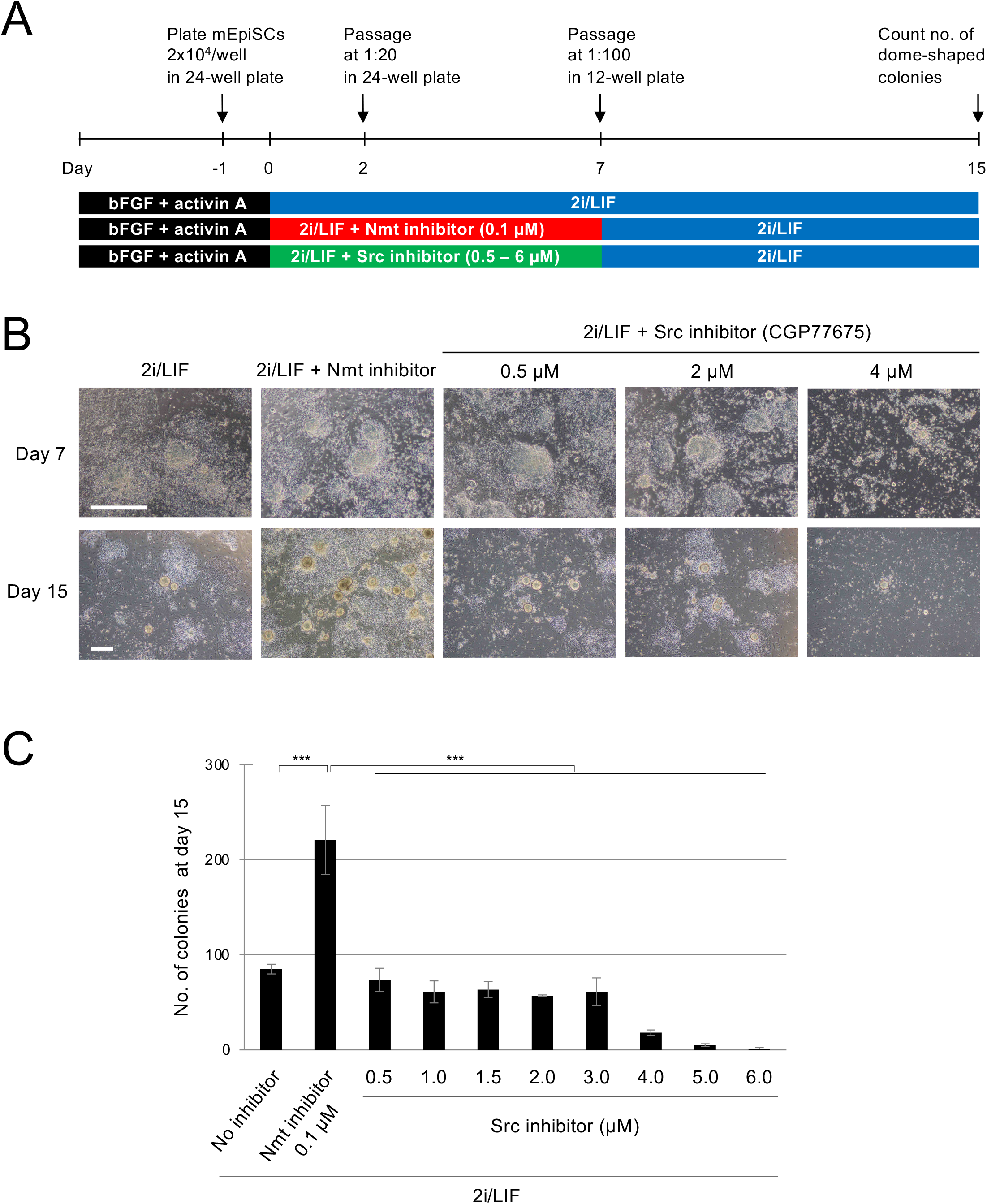
The effect of the Nmt inhibitor on the conversion of the primed to naive state is not mediated by Src signaling pathways. (A) Schematic of the protocol for the comparison of the primed-to naive-state conversion efficiency between the Nmt inhibitor DDD85464 and the Src inhibitor CGP77675. (B) A morphological view of the cells during the primed-to naive-state conversion. Scale bar: 500 µm (C) The number of dome-shaped colonies at day 15. Data are shown as mean ± SEM (*n*=3, biological replicates). Tukey-Kramer test; ^***^ *p* < 0.001.

### Inhibition of Nmt Promotes Naive State in hiPSCs

To address whether the effect of the Nmt inhibitor observed in mouse cells is generally applicable to other species, we investigated the effect of the Nmt inhibitor on human pluripotent stem cells. We initially cultured primed-state hESCs/hiPSCs in 2i medium supplemented with human LIF and the Nmt inhibitor. We then examined whether hESCs/hiPSCs are converted into the naive state as we observed in mouse cells. However, hESCs/hiPSCs differentiated gradually (Supplementary Figs. 3A and 3B), and undifferentiated cells were lost after repeated passages (Supplementary Fig. 3C). Therefore, simply adding the Nmt inhibitor to 2i/LIF does not support the naive state in human cells.

Several protocols have been reported that convert primed-state hESCs/hiPSCs into a naive state^9,10,20^. We therefore cultured naive hiPSCs in the presence of the Nmt inhibitor and investigated whether the Nmt inhibitor enhances the naive state. To quantitate the enhancement of the naive state, we utilized the EOS-GFP reporter, which is highly induced in the naive state^9^. We converted adipocyte-derived primed-state hiPSCs containing the EOS-GFP reporter into a naive state on MEF feeder cells in t2iLGö medium^20^ (Fig. 5A). We separated naive hiPSCs from MEFs by the expression of SUSD2, a naive state-specific cell surface marker^21^ (Fig. 5B). SUSD2-positive hiPSCs showed heterogeneous expression of EOS-GFP (Fig. 5C), indicating that there is a heterogeneity in the naive state under our experimental condition. This result also suggests that the EOS-GFP is a more sensitive marker of the naive state than SUSD2.

**Figure 5.**
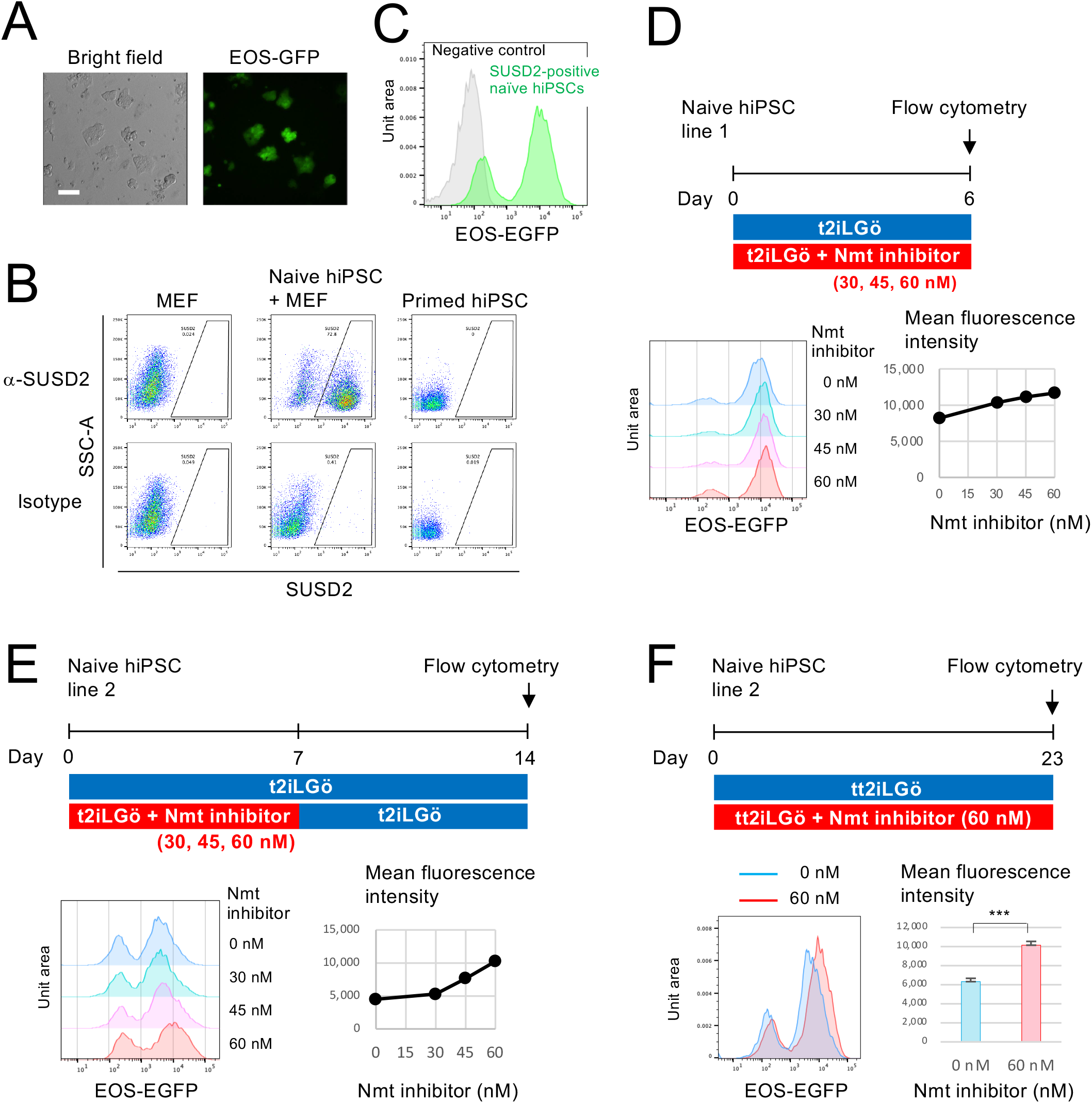
Inhibition of Nmt promotes the naive state in hiPSCs. (A) Morphology of naive hiPSCs and the expression of the EOS-GFP reporter. Scale bar: 100 µm. (B) Expression of naive-state maker SUSD2. MEF feeders could be excluded from the naive hiPSC culture by gating the expression of SUSD2. (C) The expression of EOS-GFP in the SUSD2-positive population shown in (B). Negative control is primed-state hiPSC without gating. (D, E) Dose-dependent effect of the Nmt inhibitor DDD85646 on the expression of EOS-GFP in naive hiPSCs. Note that different hiPSC lines were used in (D) and (E). (F) Long-term effect of the Nmt inhibitor on EOS-GFP expression. The mean fluorescence intensity of EOS-GFP is shown as mean ± SEM (*n*=3, biological replicates). Unpaired student’s *t*-test; ^***^ *p* < 0.001.

We initially cultured naive hiPSCs under different concentrations of the Nmt inhibitor, between 30 nM and 60 nM, for 6-days. We observed a dose-dependent increase of the expression of the EOS-GFP (Fig. 5D), suggesting that the Nmt inhibitor promoted naive state. To validate this observation, we tested the effect of the Nmt inhibitor on a different naive hiPSC line. Since we observed a decreased growth rate in the presence of the Nmt inhibitor (1:2 vs. 1:3 split with or without the 60 nM inhibitor, respectively, every 3 days), we removed the inhibitor on day 7 and continued to culture until day 14 (Fig. 5E). The growth rate recovered after removing the inhibitor, and an increase in EOS-GFP expression was observed at day 14 in a dose-dependent manner (Fig. 5E). This result is consistent with the observation in mESCs in which the effect of the Nmt inhibitor was observed in a culture without the inhibitor (Figs. 2C, 2F and 4A-4C). We also tested the effect of the Nmt inhibitor at 0.1 µM. Although this concentration was effective in mouse cells for promoting conversion to the naive state (Figs. 2C and 4A), we observed severe growth retardation in naive hiPSCs. We therefore consider 60 nM as optimal for hiPSCs.

We next examined the long-term effect of the Nmt inhibitor. We used tt2iLGö medium in which the concentration of the GSK3 inhibitor CHIR99021 was reduced from 1 µM to 0.3 µM, as this medium supports robust expansion of naive hiPSCs^20^. We set up three independent cultures with or without the Nmt inhibitor. At day 23, we quantified the expression of the EOS-GFP and confirmed that the induction of the EOS-GFP signal by the Nmt inhibitor was statistically significant (Fig 5F).

Taken together, the results indicate that suppression of Nmt enhances naive pluripotency in both mice and humans. The results also suggest the possibility that the Nmt inhibitor could be useful as a novel chemical for culturing naive-state hESCs/hiPSCs.

## DISCUSSION

In the present study, we identified Nmt as a novel target for the regulation of the naive state in both mice and humans. Nmt catalyzes the attachment of 14 carbon fatty acid myristates to the N-terminal glycine residue of proteins^12^. The significance of myristoylation during early development is underscored by the embryonic lethality observed in *Nmt1* knockout mice^18^. One of the main consequences of protein myristoylation is membrane targeting, as myristoylated proteins acquire hydrophobicity. Various signaling molecules are myristoylated, resulting in the clustering of signaling molecules at the plasma membrane and stimulation of a wide range of signaling pathways^12^. Considering this observation, an attractive model for explaining the effect of Nmt suppression on the enhancement of the naive state is the shielding of cells from external differentiation-inducing stimuli by reducing the density of signaling molecules at the plasma membrane. This concept is similar to the principle underlying the stabilization of the naive state by 2i inhibitors^13^. MEK, one of the targets of 2i, is an essential signal transducer in the FGF2-dependent differentiation pathway. Therefore, naive cells cultured in 2i medium are sequestered from a major differentiation stimulus^13^. Consistent with previous reports^17,22^, our results indicated that 2i alone was insufficient for efficiently converting from the primed to the naive state (Figs. 2F and 3G). Furthermore, the dome-shaped colony morphology, which is a characteristic feature of naive cells, was enhanced by the addition of the Nmt inhibitor to the 2i medium, indicating a non-overlapping effect between MEK and Nmt inhibitors (Fig. 1D). These observations suggest that signaling pathways other than the FGF2-MEK axis are targeted by the Nmt inhibitor.

Src is a well-characterized substrate of Nmt and is known to promote differentiation of mESCs^23,24^. Furthermore, the Src inhibitor CGP77675 stabilizes mESCs in the naive state^19^. Therefore, we initially hypothesized that Src is responsible for the conversion of the primed-state mEpiSCs to naive-state cells by Nmt suppression. However, the Src inhibitor did not enhance this conversion. This result indicated that other Nmt substrates are responsible for this conversion. Recent advances in proteomics have enabled global profiling of *N*-myristoylated proteomes, and more than 100 *N*-myristoylated proteins have been identified in HeLa cells^25^. Comparative profiling of *N*-myristoylated proteomes between naive and primed states may provide candidate *N*-myristoylated proteins regulating pluripotency.

Various roles other than plasma membrane targeting have been reported for myristoylation. A hydrophobic myristoyl moiety can alter protein folding and provide a novel interface for protein–protein interactions^26,27^. The myristoylation of proteasome components controls the shuttling of proteasome complexes between nucleus and cytoplasm, which regulates the degradation of misfolded proteins^28^. Several proteins involved in apoptosis are myristoylated following caspase-mediated cleavage, which can enhance or reduce the activity of each protein and influence the balance between cell death and survival^29^. These or other unknown mechanisms may be involved in the regulation of pluripotency observed in our study.

We observed increased expression of the naive-state marker EOS-GFP by the Nmt inhibitor. This result suggests that the use of the Nmt inhibitor may help expand the toolkit to modify the naive culture conditions for hESCs/hiPSCs. For example, ongoing attempts to generate hESC/hiPSC-derived donor organs via the formation of interspecies chimera formation^30^ requires highly competent hESCs/hiPSCs, which may be achieved by modifying the culture conditions of hESCs/hiPSCs^31^. We consider that there is room to improve the efficacy and specificity of the Nmt inhibitor because our conditional *Nmt1*-knockout experiment gave superior results compared to the Nmt inhibitor in terms of the induction level of the naive cells from primed-state cells (Figs. 2F and 3G). Recently, Nmt has attracted increasing attention as a therapeutic target in cancers, and new Nmt inhibitors are being developed accordingly^12^. The Nmt inhibitor used in the present study (DDD85646) was originally developed as a lead compound to target Nmt of *Trypanosoma brucei*^14^ and is therefore unlikely an optimal inhibitor for mammalian Nmt. The newly developed inhibitors optimized for human Nmt may regulate pluripotent stem cells more effectively.

## METHODS

### Cell Line and Cell Culture

The *Nmt1* mutant mESC clone was obtained by the gene trap method previously described^11^. The retroviral gene trap vector was inserted at the first intron of the *Nmt1* gene. The flanking sequence of the insertion site is 5’-ATCCCACGCTGGTCTCATTTGGACA-3’.

mESCs were cultured either in serum-containing medium or serum-free 2i medium depending on the purpose of the experiment. The serum-containing medium was composed of KnockOut DMEM (Cat. 10829018, Thermo Fisher Scientific) supplemented with 20% fetal bovine serum, non-essential amino acids (Cat. 11140050, Thermo Fisher Scientific), sodium pyruvate (Cat. 11360070, Thermo Fisher Scientific), 0.1 mM of 2-mercaptoethanol (Cat. M3148, Sigma) and 1,000 U/ml of leukemia inhibitory factor (LIF) (Cat. ESG1107, Merck Millipore), and mitomycin C (MMC)-treated MEFs were used as feeder cells. The serum-free 2i medium was composed of N2B27 supplemented with 1 µM of MEK inhibitor PD0325901 (Cat. Axon1408, Axon Medchem) and 3 µM of GSK3 inhibitor CHIR99021 (Cat. Axon1386, Axon Medchem). We routinely added LIF to the serum-free medium (2i/LIF), except for in the experiment shown in Fig. 1D.

mEpiSCs were cultured in DMED/F12 (Cat. 11320033, Thermo Fisher Scientific) supplemented with 20% KnockOut Serum Replacement (KSR) (Cat. 10828028, Thermo Fisher Scientific), non-essential amino acids, sodium pyruvate, 0.1 mM of 2-mercaptoethanol, 5 ng/ml of bFGF (Cat. 16100102, Katayama Chemical Industries) and 10 ng/ml of activin A (Cat. 120-14, PeproTech). MMC-treated MEFs were used as feeder cells.

Naive-state hiPSCs were established from the primed state adipocyte-derived hiPSCs^9^. We first introduced the EOS-GFP reporter vector^32^ into the primed-state hiPSCs and then induced conversion to the naive state by the protocol previously described^20^. Naive hiPSCs were maintained in t2iLGö medium, consisting of N2B27 (Ndiff227; Cat. Y40002, Takara Bio) with 1 µM of PD0325901 (Cat. 4192, Tocris), 1 µM of CHIR99021 (Cat. SML1046, Sigma-Aldrich), 10 ng/mL of recombinant human LIF (Cat. 300-05, Peprotech), and 2 µM of Gö6983 (Cat. 2285, Tocris), as previously described^9^. Naive hiPSCs were passaged every 3-5 days using Accutase (Cat. A6964, Sigma-Aldrich).

The primed-state hiPSC line MRC5iPS was generated from the human fetal lung fibroblast cell line MRC-5^33^ by retroviral transduction of reprogramming factors (Oct3/4, Sox2, Klf4, c-Myc)^7^. ALP activity was detected with VECTOR Red Alkaline Phosphatase Substrate Kit I (Cat. SK-5100, Vector Laboratories) according to the manufacturer’s instructions.

The primed-state hESC line KhES-1 was maintained as previously described^34^. To test the effect of the Nmt inhibitor DDD85646^14^ (provided by Dr. Paul Wyatt, University of Dundee, UK) on prime-state hESC, KhES-1 was passaged in 2i/LIF with or without the inhibitor and analyzed for gene expression.

For real-time PCR analysis of mESCs, MEF feeder cells were removed by plating cells on a gelatin-coated dish for 30 min during the passaging and collecting unattached cells. For real-time PCR analysis of the hESCs, MEF feeder cells were removed by separating them from clumps of hESCs under gravity sedimentation during the passaging.

### Isolation of Nmt1-revertant Clones

*Nmt1*-homozygous mESCs were transfected with pCAGGS-FLPo-IRESpuro^35^ using TransFast (Cat. E2431, Promega) to excise the *FRT*-flanked gene trap cassette. Three days after transfection, mESCs were sparsely plated on MMC-treated MEFs for single cell cloning. One week later, single cell-derived mESC colonies were picked and divided into three culture conditions: (1) with G418 (Geneticin; Cat. 10131027, Thermo Fisher Scientific), (2) with puromycin (Cat. P7255, Sigma-Aldrich), and (3) without drug selection. The parental *Nmt1*-homozygous mESC clone expresses the neomycin-resistance gene and the puromycin-resistance gene from each allele. Therefore, reversion of both alleles confers sensitivity to both G418 and puromycin.

### Immunostaining

For the immunostaining of Oct3/4 and Nanog, cells were fixed with 4% paraformaldehyde (Cat. 02890-45, Nacali Tesque) in PBS for 10 min, permeabilized with 0.2% Triton X-100 (Cat. 35501, Nacali Tesque) for 10 min, and subjected to blocking with 1% bovine serum albumin (BSA) (Cat A5611, Sigma-Aldrich) in PBS for 20 min. The following primary antibodies were used: anti-Oct3/4 mouse monoclonal antibody (1:300, clone c-10, Cat. Sc-5279, Santa Cruz Biotechnology) and anti-Nanog rabbit polyclonal antibody (1:200, Cat. RCAB002P-F, ReproCELL). Alexa Fluor 488-conjugated goat anti-mouse IgG (Cat. A-11001, Thermo Fisher Scientific) and Alexa Fluor 594-conjugated goat anti-rabbit IgG (Cat. A-11012, Thermo Fisher Scientific) were used as a secondary antibody for Oct3/4 and Nanog, respectively, and DAPI (Cat. 62248, Thermo Fisher Scientific) was used for counterstaining. For the immunostaining of SUSD2, the cells were incubated with APC-conjugated anti-SUSD2 antibody (1:20, clone W5C5, Cat. 327401, BioLegend) for 30 min in culture medium. The cells were washed three times with PBS and analyzed by FACSAria II (Becton, Dickinson and Company).

### Converting mEpiSCs to Naive-state mEpi-iPSCs

mEpiSCs (2 × 10^4^) were plated onto MMC-treated MEFs in N2B27-based medium supplemented with bFGF and activin A. The next day (day 0), the medium was changed to an N2B27-based 2i/LIF medium with or without the Nmt inhibitor (DDD85646) or the Src inhibitor (CGP77675) (Cat. 21089, Cayman Chemical). On day 2, the cells were passaged at 1:20 onto MEFs in the same medium. On day 7, the cells were passaged at 1:100 onto MEFs in 2i/LIF medium without inhibitors. On day 15, the number of dome-shaped colonies were counted.

### Generating Chimeric Mice and Determining Germline Transmission

After generating mEpi-iPSCs from mEpiSCs in serum-free 2i/LIF medium, we cultured them in serum-containing medium on MMC-treated MEFs for several days and injected them into eight cell stage embryos or blastocysts. We used ICR or BDF1-derived embryos as a host. Since the parental mEpiSCs were derived from a female 129SV mouse strain, we selected female agouti-colored chimeric mice and crossed them with male C57BL/6J mice to test germline transmission. The germline transmission was judged by the agouti coat color of the progeny.

### Generating the Conditional Allele at the *Nmt1* Locus

To conduct a conditional knockout of the *Nmt1* gene, we performed gene targeting and floxed the second exon of the *Wt* allele of the *Nmt1*-heterozygous mESC clone that we previously obtained by gene trapping^11^. The targeting vector was constructed as follows, and the PCR primer sequences are listed in Supplementary Table 1. We first PCR-amplified a genomic fragment of the first intron of the *Nmt1* gene, using primer pairs Nmt1-S-Upp1 and Nmt1-S-Low1, using the genomic DNA of the mESC line KY1.1^36^ as a template, digested with NotI and SwaI, and cloned into the NotI-SwaI site of the pMulti-Lox5171-FRT-CAG-bsd-pA-FRT (unpublished), which contains the *FRT*-flanked blasticidin S deaminase expression cassette and a single copy of the *lox5171* site at this cloning site, resulting in the pMulti-Lox5171-FRT-CAG-bsd-pA-FRT-5HR. We next amplified the *Nmt1* genomic region spanning from the first intron to the second intron, using the primers Nmt1-L-Upp2 and Nmt1-L-Low2, and the genomic region spanning from the second intron to the third intron, using the primers Nmt1-L-Upp1 and Nmt1-L-Low1. These fragments have overlapped sequences containing the *lox5171* site introduced by PCR primers. We therefore conducted fusion PCR, using the mixture of these fragments as a template, and using primers Nmt1-L-Upp1 and Nmt1-L-Low2. The fused fragments were digested with AscI and PacI and cloned into the AscI-PacI site of the pMulti-Lox5171-FRT-CAG-bsd-pA-FRT-5HR, resulting in the targeting vector pMulti-Lox5171-FRT-CAG-bsd-pA-FRT-5HR-3HR. The targeting vector was linearized with AscI, and 25 µg of the targeting vector was transfected into 1 × 10^7^ of *Nmt1*-heterozygous cells^11^ by electroporation (240 V, 500 µF) with Gene Pulser II (Bio-Rad). One week later, blasticidin S-resistant clones were picked and screened for homologous recombinants by primers Nmt1-scn1 and bsd3-1. Targeted clones were transfected with pCAGGS-FLPo-IRESpuro to remove the bsd cassette and generate the floxed *Nmt1* allele. The parental mESC line contained the *ERT2-iCre-ERT2* gene at the *Rosa26* locus^11^. Therefore, the conditional knockout of the *Nmt1* gene was achieved by treating cells with 4-hydroxytamoxifen (4HT) (Cat. H6278, Sigma-Aldrich). To confirm that the conditional allele was correctly generated, we treated mESCs with 1 µM of 4HT overnight and analyzed Cre-mediated recombination by PCR, using primers Nmt1-Flpo-Scn-F1 and Nmt1-Flpo-Scn-R1 for detecting the undeleted allele, and primers Nmt1-Flpo-Scn-F1 and Nmt1-Cre-Scn-R1 for detecting the deleted allele. We also conducted PCR by mixing all three primers in order to suppress the amplification of MEF-derived genomic DNAs that could not be eliminated by plating on a gelatin-coated dish.

### Converting mESCs to mEpiSC-like Primed-state Cells

mESCs carrying the floxed *Nmt1* allele were converted into mEpiSC-like primed-state cells according to the published protocol^17^. Briefly, mESCs were cultured in N2B27 medium supplemented with 12 ng/ml of bFGF and 20 ng/ml of activin A on a dish coated with fibronectin (Cat. 354008, Corning), and passaged at every 3–5 days. The morphology of the cell clusters became gradually flatter. We analyzed the primed-state marker *Fgf5* and the naive state marker *Dppa3* by qRT-PCR at passage nine to confirm conversion into the primed state.

### Converting mEpiSC-like Primed-state Cells to Naive-state mEpi-iPSCs by the Conditional Knockout of Nmt1

mEpiSC-like primed-state cells were plated onto MMC-treated MEFs in N2B27-based medium supplemented with 12 ng/ml of bFGF and 20 ng/ml of activin A and treated with 1 µM of 4HT for 12 hours to induce the Cre-mediated deletion of the *Nmt1* allele. The same amount of ethanol was added to the medium as a mock. The cells were maintained in the same N2B27/bFGF/activin A medium for another five days to reduce the intracellular concentration of Nmt1 protein. Then, the cells were plated onto MMC-treated MEFs in the same medium at the concentration of 2 × 10^4^ cells per well. The next day (day 0), the medium was changed to 2i/LIF medium to induce conversion to the naive state. On day 7, the cells were passaged at 1:70 onto MEFs. On day 15, the cells were stained for ALP activity, using VECTOR Red Alkaline Phosphatase Substrate Kit I (Cat. SK-5100, Vector Laboratories), according to the manufacturer’s instructions, and the number of ALP-positive colonies were counted.

### Quantitative RT-PCR (qRT-PCR)

To quantify gene expression in the mouse cells, the total RNA was extracted with RNeasy Plus Mini Kit (Cat. 74136, Qiagen) and reverse-transcribed with SuperScript III (Cat. 18080044, Thermo Fisher Scientific), using random primers (Cat. C1181, Promega). The expression levels of mRNAs encoding *Dppa3, Fgf5*, and *Actb* were quantified by real-time PCR, using the LightCycler FastStart DNA Master SYBR Green I kit (Cat. 12239264001, Roche Diagnostics) on the LightCycler (Roche Diagnostics). The primer pairs are presented in Supplementary Table 1. The amplification conditions for *Dppa3* and *Fgf5* were 95 °C for 10 min for one cycle, followed by 40 cycles of denaturation at 95 °C for 10 sec, annealing at 56 °C for 5 sec and extension at 72 °C for 20 sec. The amplification conditions for *Actb* were the same except that the annealing temperature was 55 °C. The quantity of each transcript was measured from a standard curve, and the amounts of *Dppa3* and *Fgf5* transcript were normalized to *Actb* transcript levels.

To quantify gene expression in hESCs (KhES-1), the total RNA was extracted with RNeasy Micro Kit (Cat. 74004, Qiagen) and reverse-transcribed with an RT^2^ First Strand Kit (Cat. 330404, Qiagen). The expression levels of the mRNAs were quantified using Human Embryonic Stem Cell RT^2^ Profile™ PCR Array (Cat. PAHS-081, Qiagen) and RT^2^ SYBR Green qPCR Master Mix (Cat. 330504, Qiagen). All procedures followed the manufacturer’s instructions.

### Analysis of the Localization of the myrVenus Reporter

The myrVenus reporter^15^ was cloned into the piggyBac transposon vector^37^ under the control of the CAG promoter^38^ and with the IRES-bsd selection cassette. This vector was introduced into mESCs by co-transfecting the piggyBac expression vector mPB^37^, using TransFast transfection reagent, and selected by 30 µg/ml of blasticidin S (Cat. KK-400, Kaken Pharmaceutical). We stained the plasma membrane using CellMask Deep Red Plasma Membrane Stain (Cat. C10046, Thermo Fisher Scientific) according to the manufacturer’s instructions and fixed the cells with 4% of paraformaldehyde. We captured the fluorescent images and conducted a line-plot analysis of the fluorescence signal using DeltaVision Elite (Cytiva).

### Statistical Analysis

The student’s *t*-test was conducted to compare the two groups, the Tukey-Kramer test was used for multiple comparisons between all samples, and Dunnett test for multiple comparisons was used for comparisons with a specific sample.

## AUTHOR CONTRIBUTIONS

Conceptualization, K.H. and J.T.; Methodology, J.Y. and K.H; Investigation, J.Y., H.W., K.Y., T.N., A.I., H.A., H.S., Y.T., G.K., K.H.; Writing – Original Draft, K.H.; Resources, S.O., H.N. and Y.T.; Supervision, H.A., A.U., H.S., J.T. and K.H.; Project Administration, J.T. and K.H.; Funding Acquisition, Y.T. and K.H.

## ACKNOWLEDGMENTS

We thank Dr. Paul Wyatt at the Drug Discovery Unit, School of Life Sciences, University of Dundee for providing DDD85646 (prepared with the support of Wellcome Trust grant WT 077705). We also thank Dr. Paul Tesar at Case Western Reserve University for the mEpiSCs, Dr. Kat Hadjantonakis at Sloan Kettering Institute for the myrVenus reporter vector, and Dr. Masaru Okabe for supporting generating chimeric mice. This work was supported by Grants-in-Aid for Scientific Research from the Ministry of Education, Culture, Sports, Science, and Technology of Japan (JP16H04683, JP18K19275, JP20H03174 for K.H.), JST PRESTO (K.H.), AMED (JP20bm0704035 for Y.T.), and the Cooperative Research Program (Joint Usage/Research Center program) of Institute for Frontier Life and Medical Sciences, Kyoto University (K.H.). This work was also supported in part by the research grant from the Takeda Science Foundation (K.H.), Naito Foundation (K.H.), and Daiichi Sankyo Foundation of Life Science (K.H.).

## Notes

### Competing Interest Statement

The authors have declared no competing interest.

## REFERENCES

1. Rossant J, Tam PPL. New Insights into Early Human Development: Lessons for Stem Cell Derivation and Differentiation. Cell Stem Cell 20, 18–28 (2017).

2. Evans MJ, Kaufman MH. Establishment in culture of pluripotential cells from mouse embryos. Nature 292, 154–156 (1981).

3. Martin GR. Isolation of a pluripotent cell line from early mouse embryos cultured in medium conditioned by teratocarcinoma stem cells. Proc Natl Acad Sci U S A 78, 7634–7638 (1981).

4. Tesar PJ, et al. New cell lines from mouse epiblast share defining features with human embryonic stem cells. Nature 448, 196–199 (2007).

5. Brons IG, et al. Derivation of pluripotent epiblast stem cells from mammalian embryos. Nature 448, 191–195 (2007).

6. Thomson JA, et al. Embryonic stem cell lines derived from human blastocysts. Science (New York, NY) 282, 1145–1147 (1998).

7. Takahashi K, et al. Induction of pluripotent stem cells from adult human fibroblasts by defined factors. Cell 131, 861–872 (2007).

8. Yu J, et al. Induced pluripotent stem cell lines derived from human somatic cells. Science (New York, NY) 318, 1917–1920 (2007).

9. Takashima Y, et al. Resetting transcription factor control circuitry toward ground-state pluripotency in human. Cell 158, 1254–1269 (2014).

10. Theunissen TW, et al. Systematic identification of culture conditions for induction and maintenance of naive human pluripotency. Cell Stem Cell 15, 471–487 (2014).

11. Horie K, et al. A homozygous mutant embryonic stem cell bank applicable for phenotype-driven genetic screening. Nat Methods 8, 1071–1077 (2011).

12. Yuan M, et al. N-myristoylation: from cell biology to translational medicine. Acta pharmacologica Sinica 41, 1005–1015 (2020).

13. Ying QL, et al. The ground state of embryonic stem cell self-renewal. Nature 453, 519–523 (2008).

14. Frearson JA, et al. N-myristoyltransferase inhibitors as new leads to treat sleeping sickness. Nature 464, 728–732 (2010).

15. Rhee JM, et al. In vivo imaging and differential localization of lipid-modified GFP-variant fusions in embryonic stem cells and mice. Genesis 44, 202–218 (2006).

16. Casanova E, Fehsenfeld S, Lemberger T, Shimshek DR, Sprengel R, Mantamadiotis T. ER-based double iCre fusion protein allows partial recombination in forebrain. Genesis 34, 208–214 (2002).

17. Guo G, et al. Klf4 reverts developmentally programmed restriction of ground state pluripotency. Development 136, 1063–1069 (2009).

18. Yang SH, et al. N-myristoyltransferase 1 is essential in early mouse development. J Biol Chem 280, 18990–18995 (2005).

19. Shimizu T, et al. Dual inhibition of Src and GSK3 maintains mouse embryonic stem cells, whose differentiation is mechanically regulated by Src signaling. Stem Cells 30, 1394–1404 (2012).

20. Guo G, et al. Epigenetic resetting of human pluripotency. Development 144, 2748–2763 (2017).

21. Bredenkamp N, Stirparo GG, Nichols J, Smith A, Guo G. The Cell-Surface Marker Sushi Containing Domain 2 Facilitates Establishment of Human Naive Pluripotent Stem Cells. Stem Cell Reports 12, 1212–1222 (2019).

22. Yang J, van Oosten AL, Theunissen TW, Guo G, Silva JC, Smith A. Stat3 activation is limiting for reprogramming to ground state pluripotency. Cell Stem Cell 7, 319–328 (2010).

23. Meyn MA, 3rd, Schreiner SJ, Dumitrescu TP, Nau GJ, Smithgall TE. SRC family kinase activity is required for murine embryonic stem cell growth and differentiation. Mol Pharmacol 68, 1320–1330 (2005).

24. Meyn MA, 3rd, Smithgall TE. Chemical genetics identifies c-Src as an activator of primitive ectoderm formation in murine embryonic stem cells. Sci Signal 2, ra64 (2009).

25. Thinon E, et al. Global profiling of co- and post-translationally N-myristoylated proteomes in human cells. Nat Commun 5, 4919 (2014).

26. Spassov DS, Ruiz-Saenz A, Piple A, Moasser MM. A Dimerization Function in the Intrinsically Disordered N-Terminal Region of Src. Cell Rep 25, 449–463 e444 (2018).

27. Le Roux AL, et al. A Myristoyl-Binding Site in the SH3 Domain Modulates c-Src Membrane Anchoring. iScience 12, 194–203 (2019).

28. Kimura A, Kurata Y, Nakabayashi J, Kagawa H, Hirano H. N-Myristoylation of the Rpt2 subunit of the yeast 26S proteasome is implicated in the subcellular compartment-specific protein quality control system. Journal of proteomics 130, 33–41 (2016).

29. Martin DD, et al. Rapid detection, discovery, and identification of post-translationally myristoylated proteins during apoptosis using a bio-orthogonal azidomyristate analog. FASEB J 22, 797–806 (2008).

30. Kobayashi T, et al. Generation of rat pancreas in mouse by interspecific blastocyst injection of pluripotent stem cells. Cell 142, 787–799 (2010).

31. De Los Angeles A, Pho N, Redmond DE, Jr. Generating Human Organs via Interspecies Chimera Formation: Advances and Barriers. The Yale journal of biology and medicine 91, 333–342 (2018).

32. Hotta A, et al. Isolation of human iPS cells using EOS lentiviral vectors to select for pluripotency. Nat Methods 6, 370–376 (2009).

33. Jacobs JP, Jones CM, Baille JP. Characteristics of a human diploid cell designated MRC-5. Nature 227, 168–170 (1970).

34. Suemori H, Yasuchika K, Hasegawa K, Fujioka T, Tsuneyoshi N, Nakatsuji N. Efficient establishment of human embryonic stem cell lines and long-term maintenance with stable karyotype by enzymatic bulk passage. Biochem Biophys Res Commun 345, 926–932 (2006).

35. Kranz A, et al. An improved Flp deleter mouse in C57Bl/6 based on Flpo recombinase. Genesis 48, 512–520 (2010).

36. Yagita K, et al. Development of the circadian oscillator during differentiation of mouse embryonic stem cells in vitro. Proc Natl Acad Sci U S A 107, 3846–3851 (2010).

37. Cadinanos J, Bradley A. Generation of an inducible and optimized piggyBac transposon system. Nucleic Acids Res 35, e87 (2007).

38. Niwa H, Yamamura K, Miyazaki J. Efficient selection for high-expression transfectants with a novel eukaryotic vector. Gene 108, 193–199 (1991).

