## Supplementary Information for "Inhibition of *N*-myristoyltransferase Promotes Naive Pluripotency in Mouse and Human Pluripotent Stem Cells"

### Supplementary Figure 1

Generation of chimeric mice and evaluation of germline transmission rate

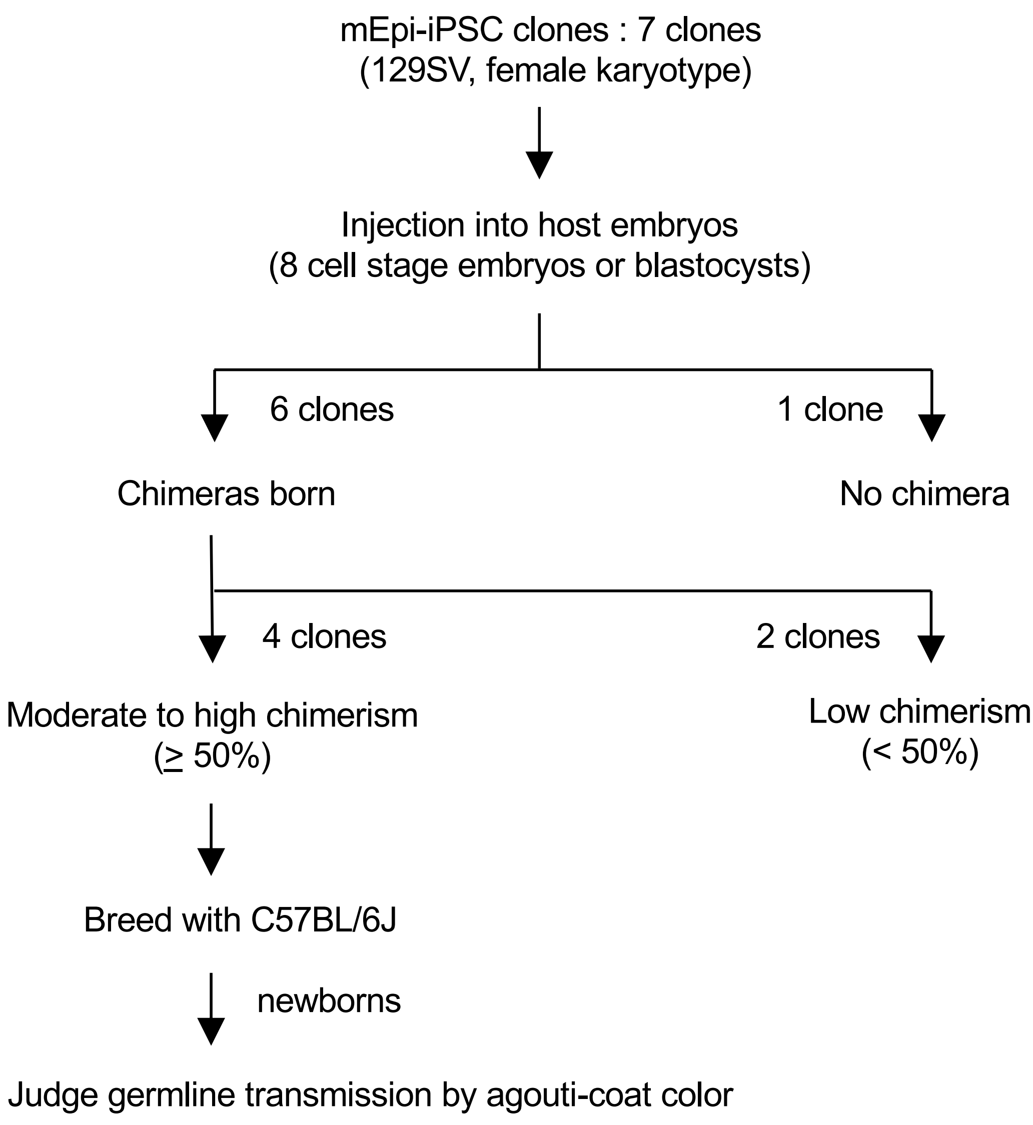


| mEpiSC-iPSC clone | Coat color of progeny agouti : black |
|---|---|
| 1 | 1:9 |
| 2 | 2:7 |
| 3 | 1:10 |
| 4 | 1:9 |

### Supplementary Figure 2

#### Generation of the conditional allele at the *Nmt1* locus by gene targeting

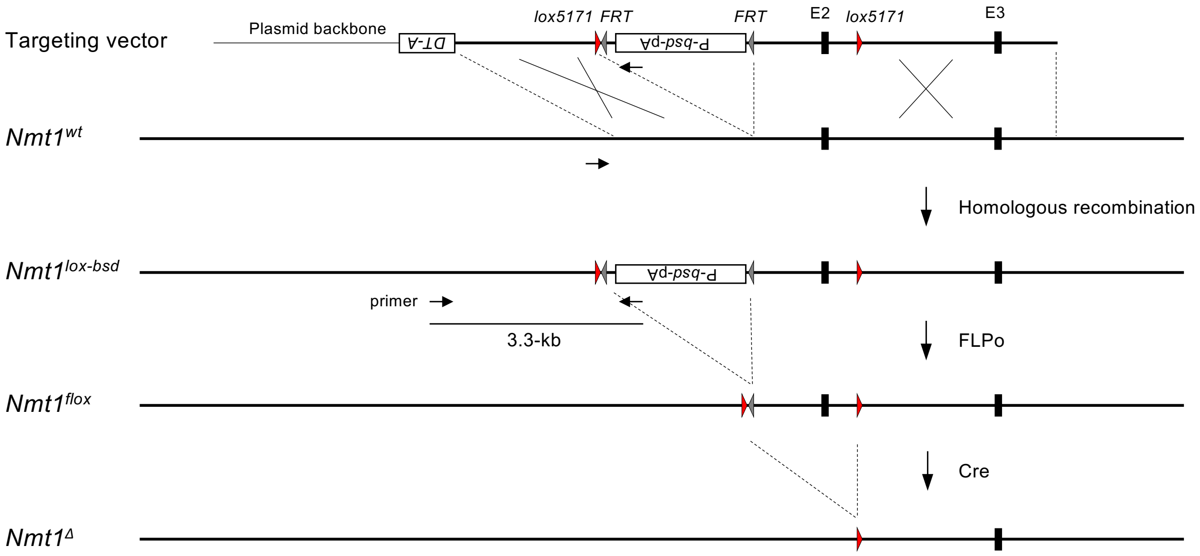


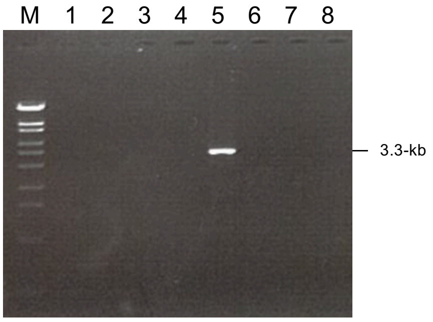


Gene targeting was conducted to the wild-type *Nmt1* allele (*Nmt1*$^{wt}$) of the heterozygous gene-trapped clone (*Nmt1*$^{m/wt}$). Homologous recombination between the targeting vector and the *Nmt1*$^{wt}$ allele was screened by PCR as shown on the left. FLPo-mediated excision of the P-*bsd*-pA cassette was identified by blasticidin S-sensitivity of the single cell-derived clones. Detection of Cre-mediated recombination was performed by 4-hydroxytamoxifen administration as shown in Figs. 3A and 3B. *DT-A*, diphtheria toxin A fragment; P, CAG promoter; *bsd,* blasticidin S deaminase gene; pA, bovine growth hormone polyadenylation signal; E, exon; M, size marker (λ/StyI digest).

### Supplementary Figure 3

Culture condition of 2i/LIF + Nmt inhibitor does not support undifferentiated state of human pluripotent stem cells

**A** ALP-staining of hiPSCs at passage 2 in 2i/LIF + Nmt inhibitor

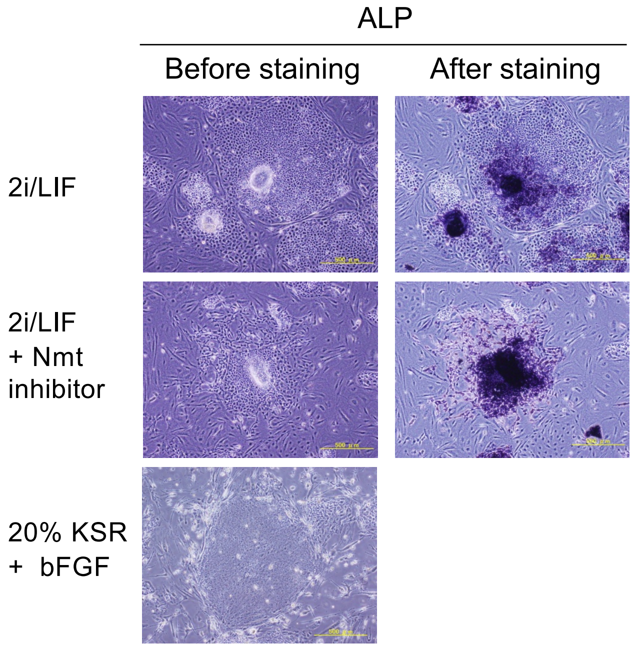


**C** Morphology of hESCs after long-term culture in 2i/LIF + Nmt inhibitor

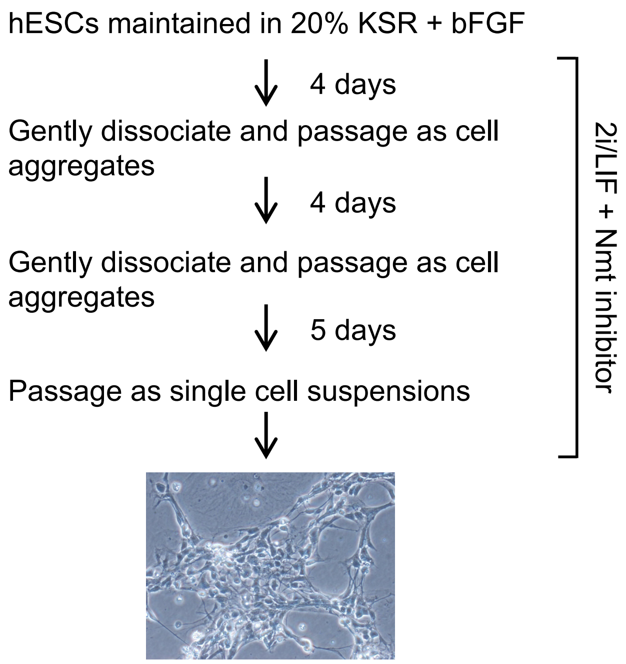


**B** Gene expression in hESCs after 4 day-culture in 2i/LIF or 2i/LIF + Nmt inhibitor relative to KSR +bFGF

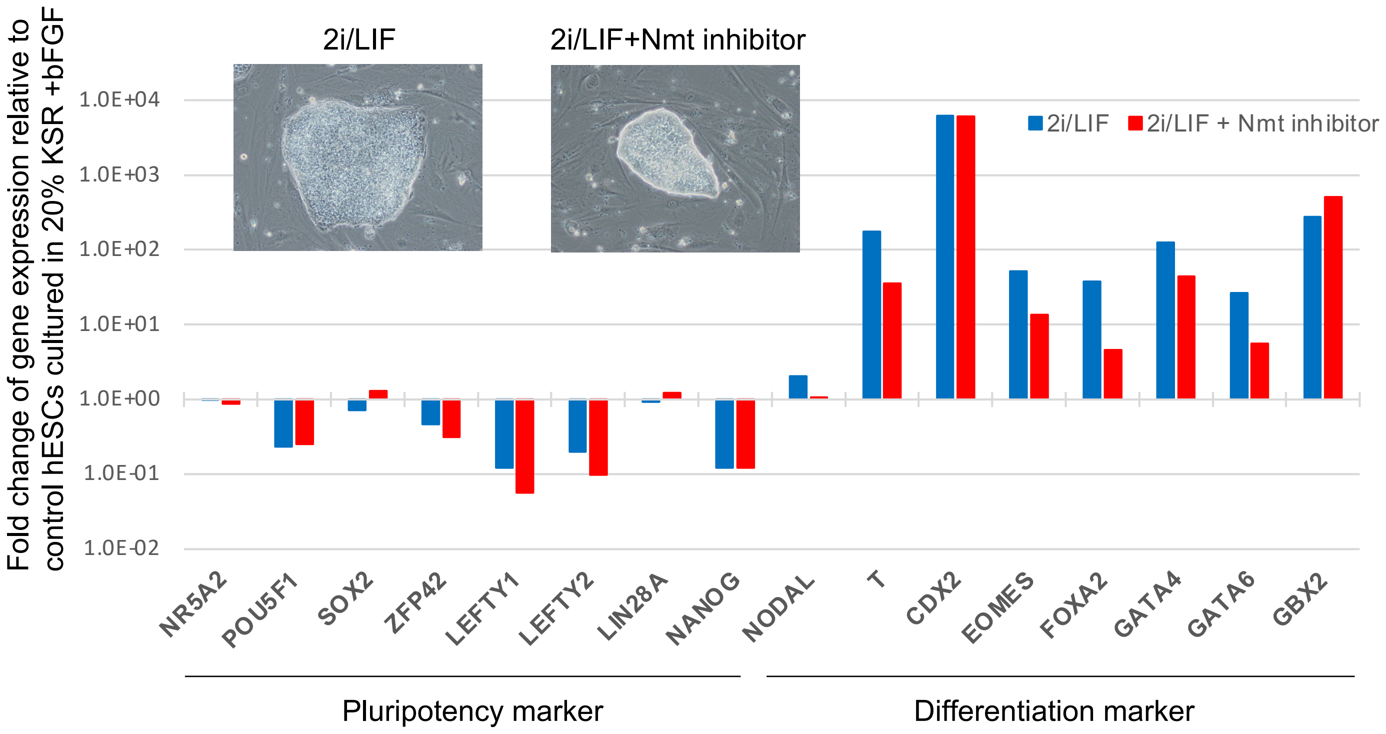

Supplementary Table 1. PCR primers.

| | |
|---|---|
| Construction of the *Nmt1* -targeting vector | |
| Nmt1-L-Upp1 | cgtttaaacggcgcgccAGCTCCAACTCCAGTACGTGCTGTCTATTC |
| Nmt1-L-Low1 | cttcgtataatgtgtactatacgaagttatGTTAACTGGTCAGGGGCTAGGTCGAGGCAC |
| Nmt1-L-Upp2 | cttcgtatagtacacattatacgaagttatTGACGAGTATTTTGACTGGTTTGCTCTTCT |
| Nmt1-L-Low2 | taccgtcgaccttaattaaATCCCACGCTGGTCTCATTTGGACATTTTA |
| Nmt1-S-Upp1 | agttctagagcggccgcCACGGCCTTGCTGTGAGAGCACGTGGGAGG |
| Nmt1-S-Low1 | cactgctcgacatttaaatGTAGCGGATGCTTCCTGCCTCCCAAGGACA |
| Screening of the *Nmt1* -targeted clones | |
| Nmt1-scn1 | GCCTCCCAAGGACAGATTCCATCTCATTCT |
| bsd3-1 | AATTGCTGCCCTCTGGTTATGTGTGGGAGG |
| Detection of the Cre/*loxP* -mediated deletion of the *Nmt1* exon 2 region | |
| Nmt1-Flpo-Scn-F1 | TTTTACATTGTCCCCACCCCAGCCT |
| Nmt1-Flpo-Scn-R1 | TTAGGCTGCTTTTCCCCCAGTAAGT |
| Nmt1-Cre-Scn-R1 | CATACTGTGCTGTCAATGTACTGTG |
| qRT-PCR | |
| mFgf5-RP-F3 | AAGTAGCGCGACGTTTTCTTC |
| mFgf5-RP-R3 | CTGGAAACTGCTATGTTCCGAG |
| Dppa3-F1 | GAGGACGCTTTGGATGATACAGACG |
| Dppa3-R1 | CAACAAAGTGCGGACCCTTCTCTTG |
| Actb-F1 | CAGGGTGTGATGGTGGGAATGGGTCAGAAG |
| Actb-R1 | TACGTACATGGCTGGGGTGTTGAAGGTCTC |
